# Increased microenvironment stiffness leads to altered aldehyde metabolism and DNA damage in mammary epithelial cells through a RhoA-dependent mechanism

**DOI:** 10.1101/2020.10.06.327726

**Authors:** Amber Wood, Heyuan Sun, Matthew Jones, Hannah Percival, Eleanor Broadberry, Egor Zindy, Craig Lawless, Charles Streuli, Joe Swift, Keith Brennan, Andrew P. Gilmore

**Author notes:** these authors contributed equally to the work (Amber Wood, Heyuan Sun, Matthew Jones, Hannah Percival). Joint senior authors (Keith Brennan, Andrew P. Gilmore). Correspondence to: Dr. Andrew P. Gilmore, Wellcome Centre for Cell-Matrix Research, Faculty of Biology, Medicine and Health, A.3034 Michael Smith Building, University of Manchester, Oxford Road, Manchester M13 9PT, Prof. Keith Brennan, Faculty of Biology, Medicine and Health, University of Manchester Oxford Road, Manchester M13 9PT.

## Abstract

Microenvironmental stiffness regulates the behaviour of both normal and cancer cells. In breast tissue, high mammographic density (HMD), which reflects greater organisation and stiffness of the periductal collagen, represents a significant risk factor for cancer. However, the mechanistic link between extracellular matrix (ECM) stiffness and increased risk of breast tumour initiation remains unclear. In particular, how increased ECM stiffness might promote genomic damage, leading to the acquisition of transforming mutations, remains to be determined. Here we determine that ECM stiffness induces changes in mammary epithelial cell (MEC) metabolism that drive genomic damage. Using a 3D-culture model, we demonstrate that genome-wide transcriptional changes in response to increased ECM stiffness impair the ability of MECs to remove reactive aldehyde species, resulting in greater accumulation of DNA damage in a RhoA-dependent manner. Together, our results provide a mechanistic link between increased ECM stiffness and the genomic damage required for breast cancer initiation.

## Introduction

Cells are exposed to a range of mechanical stimuli within their tissue microenvironment. These stimuli include properties such as the stiffness of the extracellular matrix (ECM), or cyclic strain, such as those found in lung or cardiac tissues^1 2^. Mechanotransduction describes the ability of cells to sense and convert these mechanical stimuli into biochemical signals, which mediate mechanoresponsive changes in gene expression and behaviour. Different tissues exhibit distinct mechanical properties, and cells within these tissues interpret these through integrin- and cadherin-based adhesions^3 4^. Signalling pathways coupled to these adhesion complexes then elicit intracellular responses to alter cellular behaviour. Many cellular processes, including differentiation^5^, proliferation^6^, apoptosis^7^ and migration^8^, are influenced by mechanical cues from the microenvironment. Thus, changes in the mechanical properties of a specific tissue microenvironment can have a profound effect on behaviour of the tissue and will inform our understanding of disease pathogenesis, such as fibrosis and cancer^9 10^.

Increased stiffness of the stromal ECM can increase mammary tumour progression and invasion, through alterations in mechanosignalling^11^. However, altered mechanosignalling may also be linked to breast cancer initiation. After age, total breast mammographic density - the area of radio-opaque fibroglandular tissue seen on a mammogram - is the second largest independent risk factor for breast cancer^12^. High mammographic density (HMD) is associated with changes in the composition and organisation of the collagen within the periductal stroma, leading to increased stromal stiffness^13^. Thus, altered mechanosignalling may promote changes within breast cell behaviour that promote tumour initiation^14^.

Here we asked whether changes in mechanosignalling within mammary epithelial cells can drive the phenotypic changes required to promote genomic damage and cancer initiation. We used a 3D hydrogel system composed of an interpenetrating network of reconstituted basement membrane and alginate which can be mechanically tuned to mimic variations in the mechanical tissue microenvironment^15^. Mammary acini within a stiff micro-environment developed a pre-malignant phenotype characterised by excessive and irregular growth, a loss of differentiation and tissue specific gene expression, and an increased ability to grow in soft agar. A comparison of gene expression between cells grown in a soft or stiff 3D-ECM highlighted widespread changes in metabolic pathways. In stiff matrices, we identified decreased expression of several isoforms of aldehyde dehydrogenase, the consequence of which was increased reactive aldehydes and the accumulation of DNA damage. These changes were mediated through Rho-dependent mechanosignalling. Together these results establish a link between ECM stiffness and the induction of genomic damage and a breast cancer-like phenotype through altered metabolism of reactive aldehydes.

## Results

### Increased ECM stiffness inhibits differentiation of mammary epithelial cells

Both normal and transformed mammary epithelial cells have been shown to undergo phenotypic changes in response to alterations in the mechanical stiffness of their microenvironments^10,15,16^. To understand how differences in ECM stiffness-driven mechanosignalling might promote early events in cancer initiation, we first asked how increased ECM stiffness affects the differentiation state of murine mammary epithelial cells (mMECs). We utilised a previously described 3D-culture model composed of interpenetrating networks (IPNs) of reconstituted basement membrane (Matrigel) and alginate. These gels are mechanically tuneable through addition of calcium sulphate (CaSO_4_)^15^ (Fig. S1a). mMECs freshly isolated from pregnant mice were embedded in Matrigel/alginate IPNs, with CaSO_4_ concentrations of either 0 mM or 2.4 mM to create soft and stiff conditions. mMECs were cultured in the 3D IPNs for 7 days, treated with prolactin to induce milk production, and then extracted and immunostained for β-casein (Fig. 1a). The number of β-casein positive cells was also quantified. mMECs cultured within a stiffer 3D ECM showed significantly less staining for β-casein, indicating impaired differentiation.

**Figure 1.**
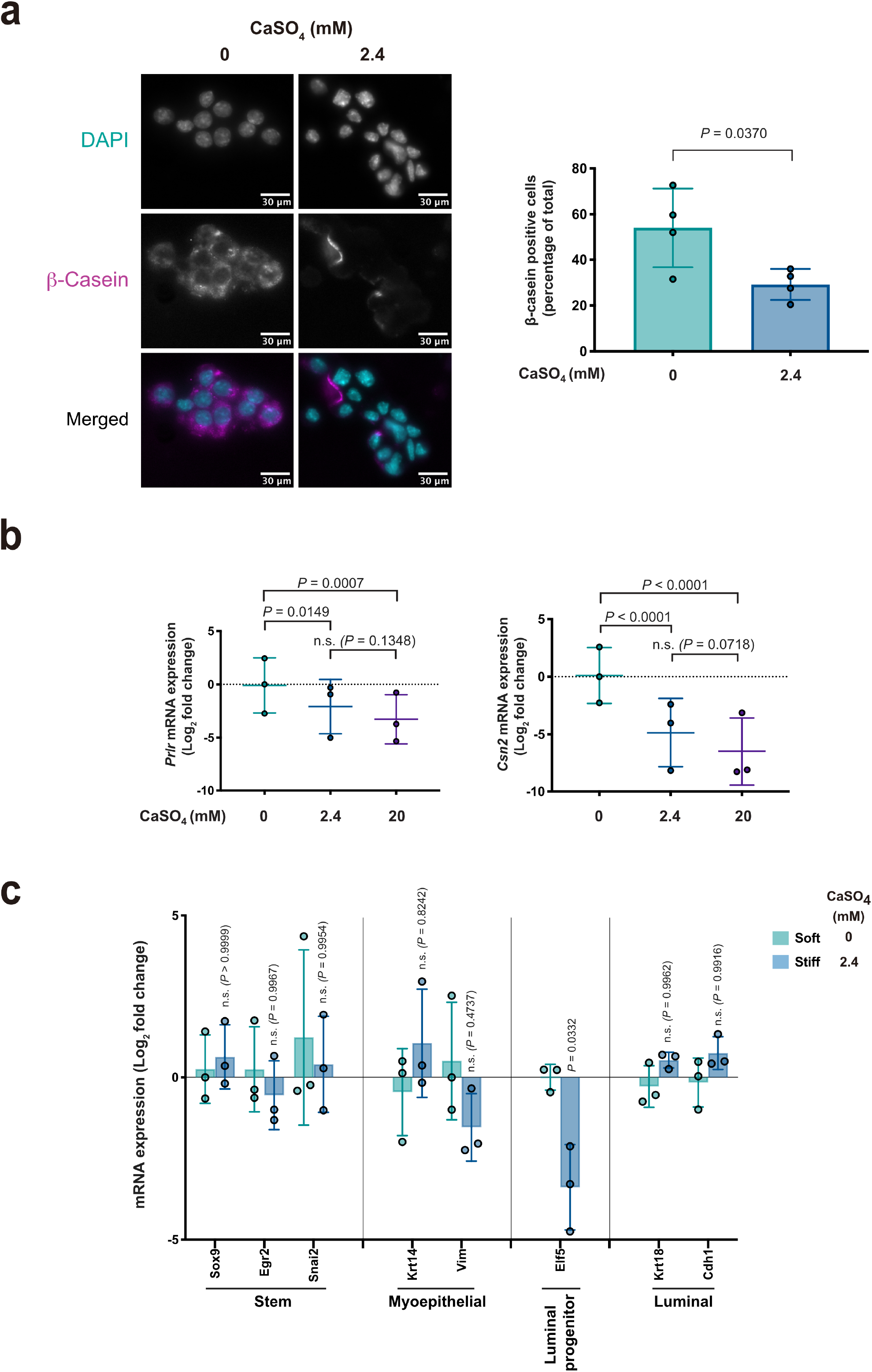
Increased ECM stiffness inhibits differentiation of mammary epithelial cells. **(a)** Representative immunofluorescence (IF) images of mMECs stained for β-casein following culture in 3D gels of different stiffnesses, (scale bars, 30μm), (left panel); and accompanying quantification (right panel). Data points shown represent results from *n* = 4 biological replicates per condition; bar represents mean ± SD. Two-tailed Student’s *t*-test, (*t* = 2.67, df = 6). **(b)** Log_2_ fold-change in gene expression of *Prlr* (left), and *Csn2* (right), normalised to *Gapdh*, as determined by RT-qPCR in EpH4 cells, relative to the 0 mM CaSO_4_ condition. Mean ± SD, *n* = 3 per condition, across independent experiments (represented by data points). Twoway ANOVA with Tukey’s post-hoc test, (F(2,9) = 16.7024, and F(2,9) = 57.2581 for *Prlr* and *Csn2*, respectively). **(c)** Log_2_ fold-change in gene expression of mammary cell lineage markers as determined by RT-qPCR in EpH4 cells, relative to the 0 mM CaSO_4_ condition. Mean ± SD, n = 3 for each condition across independent experiments (represented by data points). Two-way ANOVA with Tukey’s post-hoc test, (F(1,36) = 2.7866).

We next assessed the mechanosensitive response of EpH4 cells within 3D IPNs. EpH4s are a non-transformed murine luminal epithelial cell line^17^. Like primary mMECs, EpH4 cells are able to form 3D acini within Matrigel and differentiate to produce milk proteins in response to prolactin. We embedded single cell suspensions of EpH4 cells within IPNs with CaSO_4_ concentrations of 0 mM, 2.4 mM, and 20 mM, generating gels of increasing mechanical stiffness (Fig. S1a). Following 10 days of culture in this model, we quantified EpH4 acinar size. Acinar area was significantly larger within IPNs with 2.4mM CaSO_4_ and they failed to hollow. There was no further increase in acinar size with 20mM CaSO_4_ (Fig. S1b).

To assess the effect of ECM stiffness on EpH4 cell differentiation, we carried out quantitative reverse transcriptase PCR (RT-qPCR) to measure relative expression of prolactin receptor (*Prlr*), and beta-casein (*Csn2*), two markers of milk production. Expression of both genes was significantly reduced in stiffer 3D ECM compared with soft. As with acinar size, IPNs with 2.4 mM CaSO_4_ significantly reduced expression of *Prlr* and *Csn2*, and there was no further reduction in expression when cells were grown in the stiffest ECM (Fig 1b). Based on this, all further experiments used only the addition of 2.4 mM CaSO_4_ for stiff 3D IPNs and compared these with soft gels where no additional CaSO_4_ was included.

Breast cancer is often associated with reversion of MECs to a more stem-like state. As we saw a mechanosensitive change in MEC-specific genes involved in milk expression, we asked whether there was a linked change in cell fate. We used RT-qPCR analysis of EpH4 cells grown in 3D soft and stiff (2.4 mM CaSO_4_) to compare marker gene expression for mammary stem cells, myoepithelial cells, luminal progenitors and differentiated luminal epithelial cells (Fig. 1c). There were no significant changes in the expression of the mammary stem cell markers *Sox9, Egr2*, or *Snai2*. Similarly, there was no evidence of changes in expression of marker genes for myoepithelial cells (*Krt14* and *Vim*), or differentiated luminal epithelial cells (*Krt18* and *Cdh1*). The luminal progenitor marker *Elf5* did show a reduced expression in stiff conditions relative to soft, but overall these data did not indicate a shift towards a less-differentiated cell type or a cell type that cannot express milk.

Together, these results suggest that increased ECM stiffness disrupts the terminal differentiation of EpH4 cells and primary mMECs.

### Increased ECM stiffness alters global gene expression in mammary epithelial cells

To understand how altered ECM stiffness might impact on mammary epithelial cell behaviour and differentiation, we performed an unbiased global transcriptome analysis using RNA sequencing (RNAseq). We compared gene expression across the transcriptome in EpH4 cells grown in either soft or stiff 3D IPNs. Overall, approximately 1500 genes were differentially expressed between EpH4 cells cultured in soft and stiff 3D culture conditions (Fig. 2a-b). The majority of these genes showed reduced expression in response to increased ECM stiffness. We specifically looked within the RNAseq data for genes involved in milk production. In agreement with the RT-qPCR analysis, expression of several genes associated with milk production, including a number of casein genes and the prolactin receptor, were significantly downregulated in the RNAseq data from cells within the stiff IPNs, compared to those in soft conditions (Fig. 2c). Many studies on mechanosignalling have examined cells cultured on 2D substrates of different stiffnesses. Interestingly, the differences in gene expression seen in Eph4 cells were unique to cells grown in 3D matrices as they were not observed when cells were grown on 2D soft and stiff substrates (Fig. S2a-b). Indeed, there were more significant differences in gene expression between cells grown in 2D compared to 3D than between cells grown in soft compared to stiff environments.

**Figure 2.**
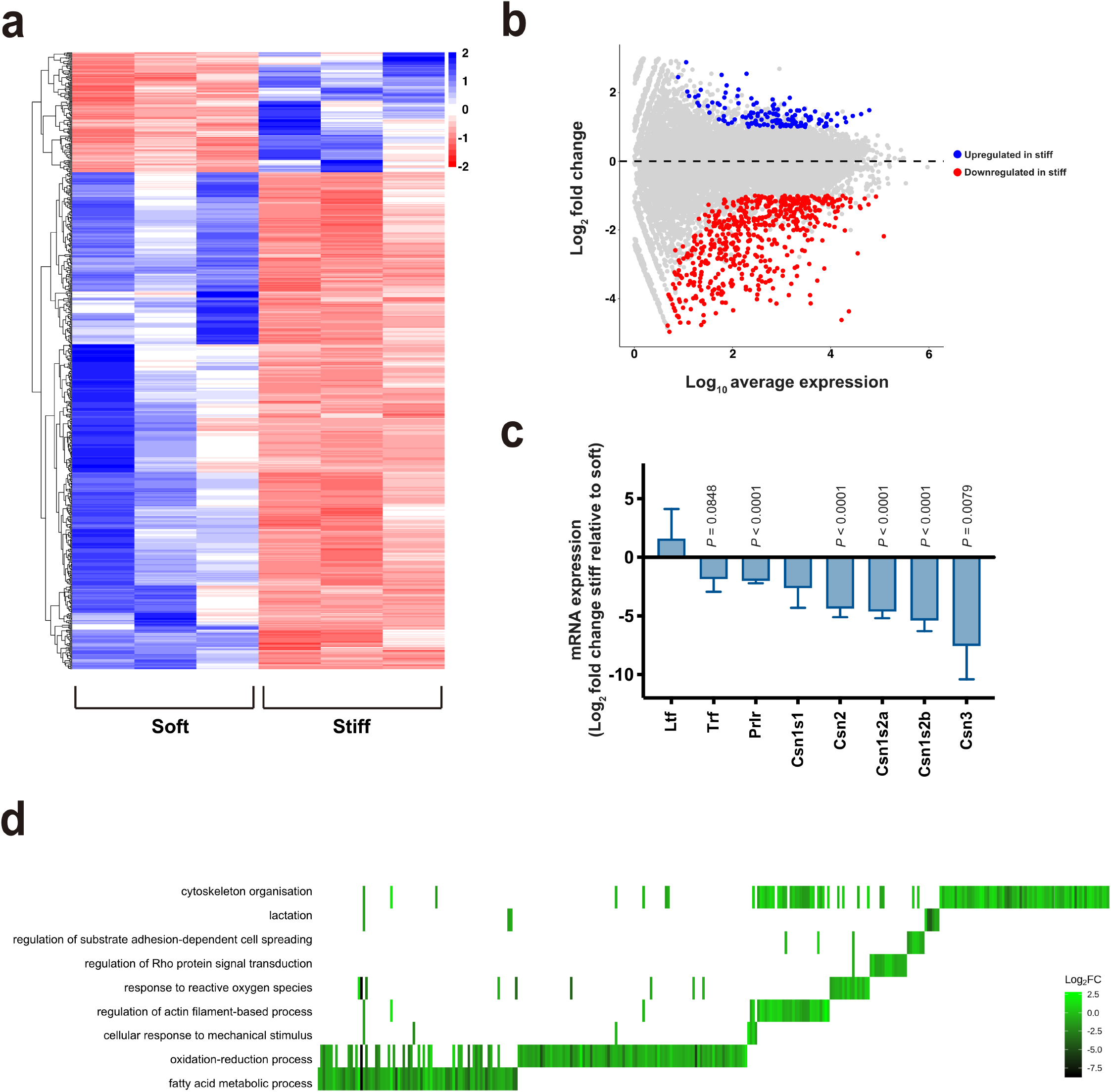
Increased ECM stiffness alters global gene expression in mammary epithelial cells. **(a)** Hierarchical clustering heatmap representing significantly differentially expressed genes in EpH4 cells cultured in 3D gels of different stiffnesses, as determined by RNAseq. Three independent biological replicates of both soft and stiff 3D cultures are shown. **(b)** MA plot generated from RNAseq data, showing genes that are significantly upregulated (blue) and downregulated (red) in EpH4 cells grown in the stiff condition, relative to soft. Data shown are the mean from the 3 biological replicates in (a). **(c)** Log_2_ fold-change in expression of genes associated with milk production in EpH4 cells grown in the stiff condition relative to soft, as determined by RNAseq. Error bars represent SE (*n* = 3), statistical significance was determined using DESeq2. **(d)** Heatplot showing significant GO terms associated with differentially expressed genes, and their Log_2_ fold-change values in the stiff condition, relative to soft.

Taken together, these results highlight the impact of altered mechanosignalling on global gene expression in non-transformed mammary epithelial cells within a 3D ECM.

### ECM stiffness-induced downregulation of aldehyde dehydrogenases impairs the ability of mammary epithelial cells to oxidise reactive aldehydes

To obtain an insight into how ECM stiffness might promote the initiation of tumorigenesis, we asked which biological processes were most influenced by the gene expression changes identified. We performed gene ontology enrichment analysis on the RNAseq dataset (Fig 2d). As well as lactation, we saw expected changes in pathways associated with cell adhesion, cytoskeletal organisation, and mechanosensing, such as those associated with Rho GTPase signalling. However, the GO enrichment analysis also identified several other key pathways that showed specific differences between EpH4 cells in the 3D soft and stiff cultures. Interestingly, some of the largest expression changes were observed in genes relating to a number of metabolic processes. These included fatty acid metabolism, response to reactive oxygen species, and oxidation/reduction processes. As other recent studies have identified that metabolic processes are regulated by changes in mechanosignalling^18,19^, we speculated that changes in these processes might be linked to increased tumour initiation.

To investigate this further, we looked through the RNAseq dataset to identify the molecular function terms associated with differentially expressed genes (Fig. 3a). Some of the most prominent changes were linked with oxidoreductase activity, in particular the oxidation of aldehydes. Amongst the largest expression differences observed between EpH4 cells in soft and stiff IPNs was that of genes coding for aldehyde dehydrogenases (ALDHs). Seven of the ALDH genes were significantly downregulated in stiff ECM compared to soft (Fig. 3b). Aldehydes are highly reactive intermediary products of many metabolic pathways, and represent a major source of endogenous DNA damage^20^. As such, any changes in a cell’s ability to oxidise aldehydes might lead to their accumulation, resulting in subsequent DNA damage and impaired protein function through formation of adducts. Interestingly, *Aldh1a1*, the isoform frequently used as a marker of breast cancer stem cells^21^, was not significantly altered between the soft and stiff conditions (Fig. 3b), in agreement with our observation that there was no shift in fate within the mammary epithelial cell lineage (Fig. 1c).

**Figure 3.**
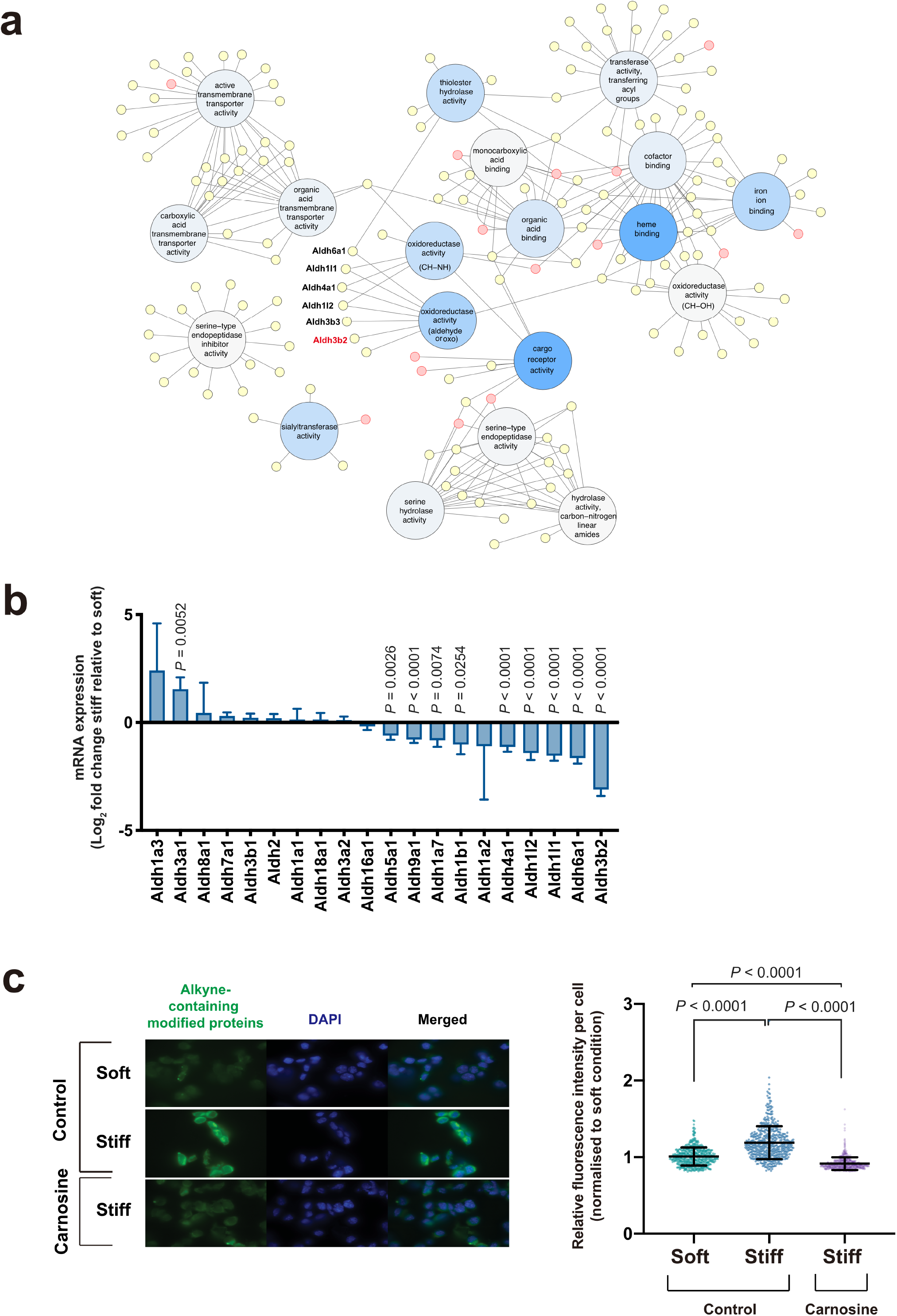
ECM stiffness-induced downregulation of aldehyde dehydrogenases impairs the ability of mammary epithelial cells to oxidise reactive aldehydes. **(a)** Bipartite graph showing significant GO molecular function terms (large circles) and their associated differentially expressed genes (small circles). Red circles represent genes associated with the “Neoplasms” MeSH term. The text indicates the six ALDH isozymes associated with oxidoreductase activity showing significant downregulation in EpH4 cells grown in stiff relative to soft 3D ECM. **(b)** Log_2_ fold-change in expression of ALDH isozymes in EpH4 cells grown in the stiff 3D ECM relative to soft, as determined by RNAseq. Error bars represent SE (*n* = 3), and statistical significance was determined using DESeq2. **(c)** Representative images shown in the left panel of EpH4 cells grown in soft or stiff 3D ECM, with or without addition of carnosine, following treatment and staining with Click-iT™ Lipid Peroxidation Imaging Kit. Right panel shows single cell IF quantification. Mean ± SD, (*n* = 510 cells/condition, across two independent experiments). Data points represent the fluorescence intensity of individual cells, normalised to the median value for the soft, untreated condition. Kruskal-Wallis test with Dunn’s multiple comparisons post-hoc test.

As ALDHs catalyse the oxidation of aldehydes, we sought to determine whether their downregulation in cells grown within stiff ECM resulted in accumulation of reactive aldehydes. To test this, we cultured EpH4 cells in soft and stiff ECM for 24 hours prior to incubation with alkyne-modified linoleic acid (LAA). LAA incorporates into cell membranes, where it can be oxidised during lipid peroxidation. Lipid peroxidation is one of the major sources of endogenous aldehydes. When LAA is oxidised to hydroperoxides, these subsequently decompose to aldehydes which can modify proteins at nucleophilic side chains, leaving an alkyne group that can be detected by Click-iT fluorescent staining. EpH4 cells cultured in soft and stiff 3D IPNs were subsequently isolated, dissociated into single cells and stained with Alexa Fluor 488 Azide. Fluorescence intensity was quantified for single cells within each population. There were significantly higher levels of alkyne-modified proteins in cells isolated from stiff 3D IPNs compared to those from soft conditions (Fig. 3c). The higher level of protein modification in stiff gels could be rescued by the addition of carnosine, a dipeptide that scavenges unsaturated aldehydic lipid oxidation products. These results show that increased ECM stiffness results in downregulated expression of ALDH genes, with resulting impairment of the oxidisation and removal of reactive aldehydes.

### Increased ECM stiffness induces DNA damage in mammary epithelial cells via downregulation of aldehyde dehydrogenases

As we observed an accumulation of reactive aldehydes in EpH4 cells cultured in a stiffer ECM, we next asked whether these cells were subject to higher levels of DNA damage compared to cells in soft. To assess DNA damage, we visualised phosphorylation of histone H2AX at Ser 139 (γ-H2AX), which is a sensitive marker of double-stranded DNA breaks^22^. EpH4 cells were seeded into soft or stiff ECM and cultured for 24-hours prior to extraction, dissociated into single cells and immunostained for γ-H2AX (Fig. 4a). We quantified the number of γ-H2AX foci in single cells (Fig. 4b). There were significantly more γ-H2AX foci in cells cultured within stiff ECM compared to those cultured in soft, indicating that they had acquired more DNA damage. We confirmed these results by immunostaining cells with antibodies to phosphorylated CHK1 (Ser345) and CHK2 (Thr68), as downstream markers of ATM and ATR activation following DNA damage (Fig. S4a). Both CHK1 and CHK2 showed increased phosphorylation in cells isolated from the stiff ECM compared with soft.

**Figure 4.**
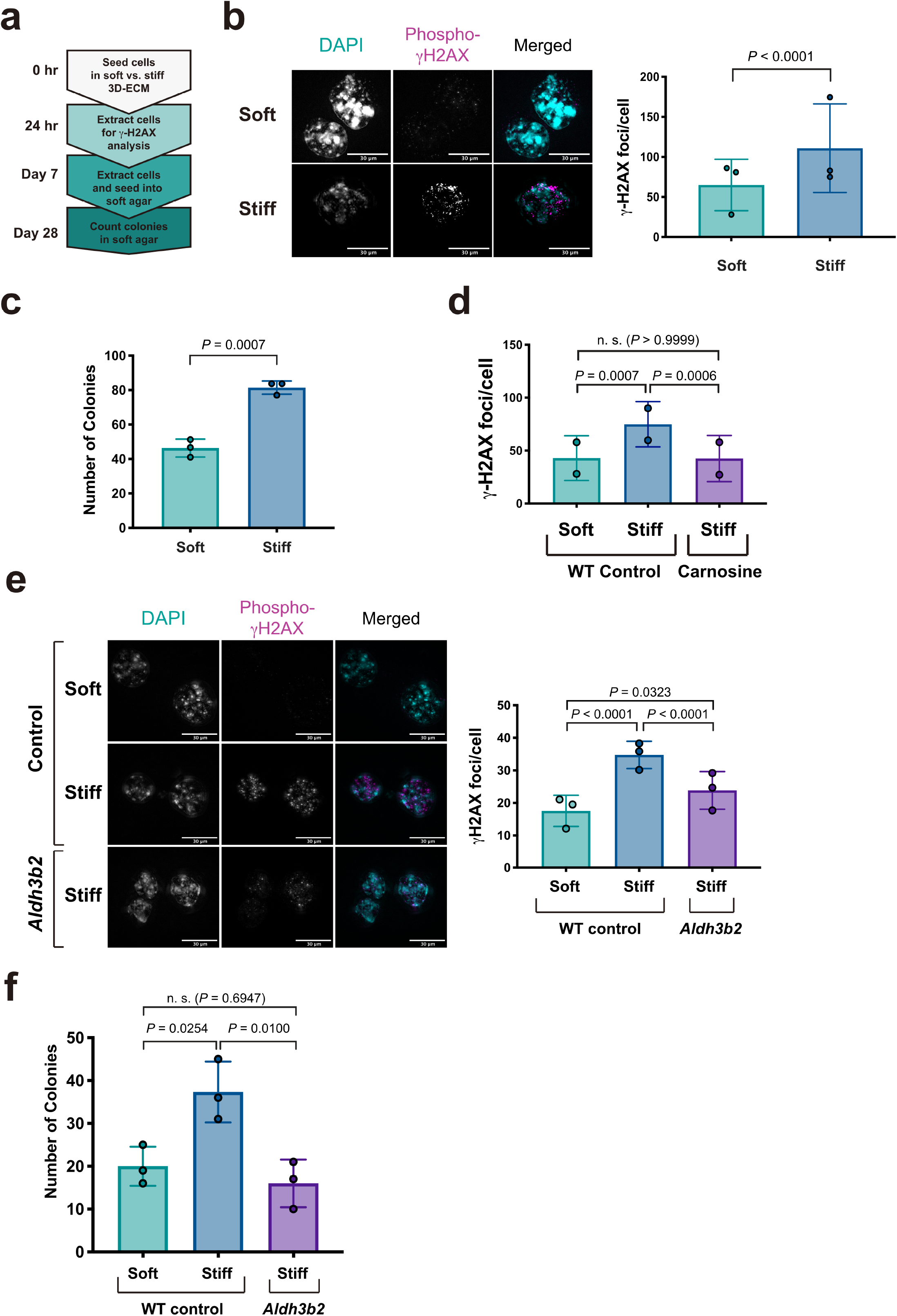
Increased ECM stiffness induces DNA damage in mammary epithelial cells via downregulation of aldehyde dehydrogenases. **(a)** Schematic of experimental design. Cells were cultured in either soft or stiff 3D ECM for either 24 hrs or 7 days. At each time point, cells were extracted and dissociated into single cells prior to either immunostaining or seeding into soft agar. **(b)** Left panel shows representative IF images of EpH4 cells immunostained for phospho-γH2AX (Ser139) following culture in 3D gels of different stiffnesses for 24 hrs (scale bars, 30 μm). Right panel shows accompanying quantification of number of phospho-γH2AX (Ser139) foci/cell. Mean ± SD, *n* = 3 per condition from independent experiments. Data points represent mean number of foci for each independent experiment, calculated from 40-50 cells/condition. Two-way ANOVA, (F(1,284) = 47.6949). **(c)** Number of colonies formed in soft agar following culture in soft or stiff 3D ECM for seven days. Colonies were quantified after 21 days in soft agar. Mean ± SD, *n* = 3 per condition from independent experiments, each performed in triplicate. Data points represent mean number of colonies from triplicates for each independent experiment. Two-tailed Student’s *t*-test, (*t* = 9.4297, df = 4). **(d)** Quantification of phospho-γH2AX foci in EpH4 cells following culture in soft or stiff 3D ECM with or without carnosine. Mean ± SD, *n* = 2 per condition from independent experiments. Data points represent mean number of foci for each independent experiment, calculated from 20-55 cells/condition. Two-way ANOVA with Tukey’s post-hoc test, (F (2,228) = 9.4557). **(e)** WT EpH4 cells or those stably overexpressing *Aldh3b2* grown in soft or stiff 3D ECM as indicated. Upper panel shows representative IF images of EpH4 cells stained for phospho-γH2AX (Ser139), (scale bars, 30μm). Lower panel - quantification of phospho-γH2AX (Ser139) foci/cell. Mean ± SD, *n* = 3 per condition from independent experiments. Data points represent mean number of foci for each independent experiment, calculated from 30 cells/condition. Two-way ANOVA with Tukey’s post-hoc test, (F (2,261) = 24.359431). **(f)** Number of colonies formed in soft agar following culture in soft or stiff 3D ECM for seven days. Mean ± SD, *n* = 3 per condition from independent experiments, each performed in triplicate. Data points represent mean number of colonies from triplicates for each independent experiment. One-way ANOVA with Tukey’s post-hoc test, (F(2, 6) = 11.3094).

If EpH4 cells suffered more genomic damage in stiff 3D cultures, then we would predict that these cells would be more prone to transformation. To test this, we used the soft agar colony formation assay to assess anchorage-independent growth, an established marker of transformation^23^. EpH4 cells were cultured in soft or stiff 3D IPNs for 7 days prior to extraction, dissociation into single cells, and re-seeding into soft agar for a further 21 days. The number of colonies formed was then quantified. Cells cultured for one week in stiff ECM formed more colonies in soft agar than those cultured in soft ECM (Fig. 4c).

We asked if the increase in reactive aldehydes in cells in stiff ECM might cause the DNA damage and colony formation in soft agar. First, we treated cells in stiff ECM with carnosine, and, following isolation of single cells, we compared the number of γ-H2AX foci with that in untreated cells, or cells in soft ECM. Treatment with carnosine reduced the number of γ-H2AX foci in cells grown in stiff ECM to levels similar to those seen in cells grown in soft ECM (Fig. 4d). We next determined whether the down regulation of ALDH genes in stiff ECM could be directly linked to genomic damage. We generated EpH4 cells which stably expressed *Aldh3b2* under a constitutive EF1α promoter, via lentiviral transduction. We chose *Aldh3b2* as this was the most significantly downregulated ALDH isoform in the RNAseq data. When cultured in stiff ECM, *Aldh3b2*-expressing EpH4 cells accumulated less DNA damage than wildtype (WT) cells, as quantified by the number of γ-H2AX foci, and the formation of fewer colonies when subsequently cultured in soft agar, (Fig. 4e-f).

It was possible that increased genomic damage could have resulted from suppression of DNA repair pathways. To determine if impairment of these pathways contributed to the increased γ-H2AX foci following culture in stiff IPNs, we used our RNAseq dataset to ascertain the fold change in expression of genes involved in DNA repair between the two conditions. There was no reduced expression of key DNA repair genes in EpH4 cells grown in stiff compared to soft 3D IPNs (Fig S3a-e). Instead, most DNA repair genes showed an increase in expression in the stiffer cultures, although this was mostly not at the level of significance. However, this would be in accordance with cells accumulating greater DNA damage, showing that the cells had not lost their ability to respond to DNA damage.

Together, these data indicate that culture of MECs within a stiffer ECM results in downregulation of ALDH genes, increased accumulation of reactive aldehydes, leading to increased DNA damage and transformation.

### ECM stiffness-induced YAP activation affects acini morphology, but does not mediate stiffness-induced DNA damage

We next sought to determine the signalling mechanisms by which mechanical stiffness contributed to increased DNA damage. The YAP/TAZ pathway is an established mechanosensitive signalling pathway with known links to proliferation and tumourigenesis^24^. Our RNAseq dataset identified upregulation of several YAP/TAZ target genes when EpH4 cells were cultured in the stiff ECM, indicating that the pathway is activated (Fig. 5a). To examine the role of YAP signalling, we generated two stable EpH4 cell lines by lentiviral transduction. In one we expressed YAP-4SA, a variant of YAP with four inhibitory serine phosphorylation sites substituted to alanine, resulting in its constitutive activation^25^. In the other we expressed a dominant negative TEAD2 (TEAD2dn) which lacks the DNA binding domain, previously shown to block YAP-dependent transactivation^26^. We then assessed the effect of activating YAP signalling in soft ECM and inhibiting it in stiff.

**Figure 5.**
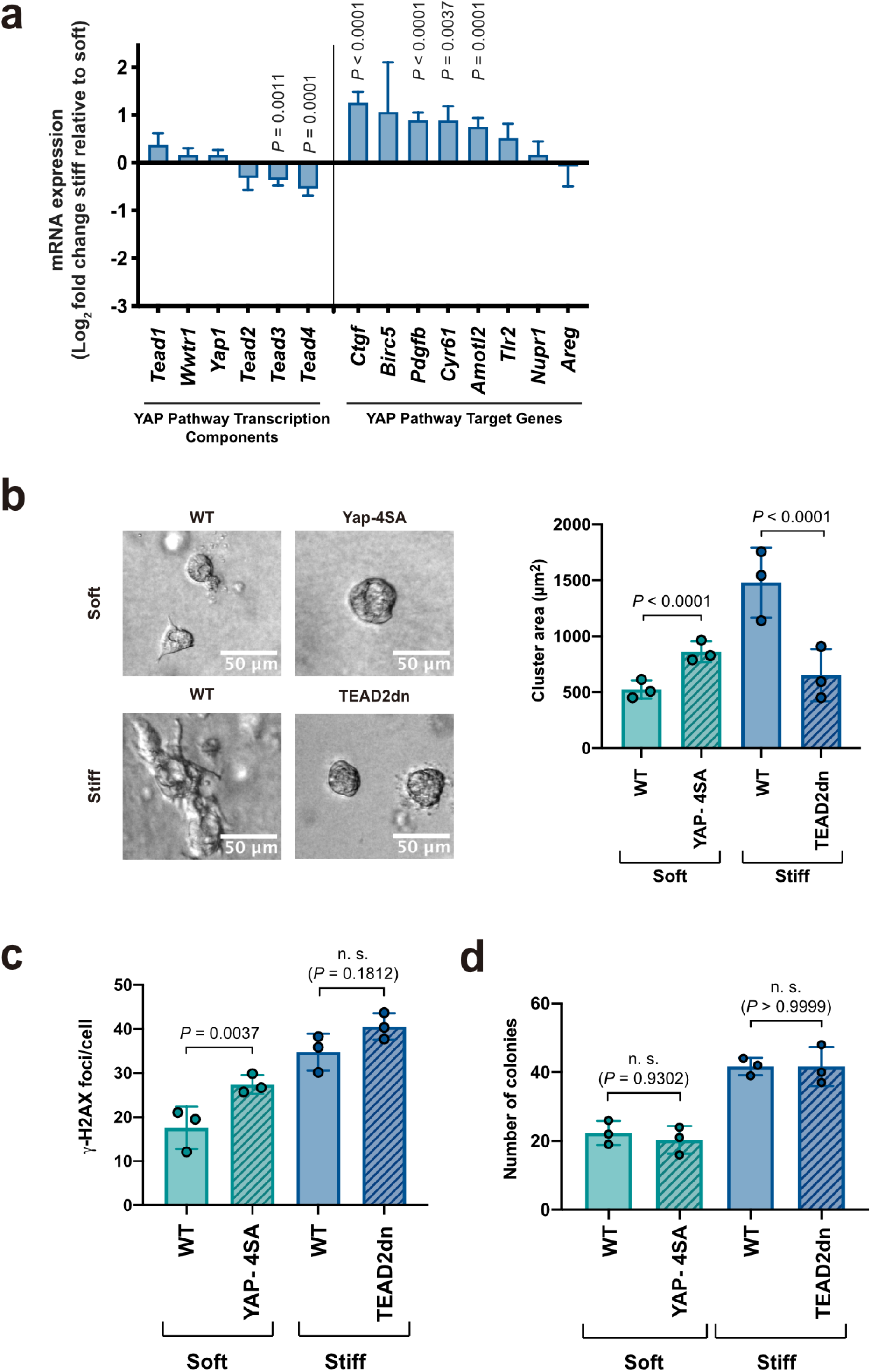
ECM stiffness-induced YAP activation affects acini morphology, but does not mediate stiffness-induced DNA damage. **(a)** Log_2_ fold-change in expression of genes involved in the YAP/TAZ signalling pathway in EpH4 cells grown in the stiff condition relative to soft, as determined by RNAseq. Error bars represent SE (*n* = 3). Statistical significance was determined using DESeq2. **(b)** Left panel shows representative brightfield images of EpH4 acini, WT, or those stably expressing YAP-4SA or TEAD2dn, following 10 days of culture in gels of different stiffnesses (scale bars, 50μm). Right panel shows accompanying quantification. Mean ± SD, *n* = 3 per condition from independent experiments. Data points represent mean acinar area for each independent experiment, calculated from 45-105 acini/condition. Two-way ANOVA with Tukey’s post-hoc test, (F (3,937) = 200.3521). **(c)** Quantification of phospho-γH2AX (Ser139) foci in EpH4 cells, WT or stably expressing YAP-4SA or TEAD2dn, following 24 hours of 3D culture. Mean ± SD, *n* = 3 per condition from independent experiments. Data points represent mean number of foci for each independent experiment, calculated from 30 cells/condition. Two-way ANOVA with Tukey’s post-hoc test, (F (3,348) = 23.9198). **(d)** Number of colonies formed in soft agar following culture in soft or stiff 3D ECM for seven days. Mean ± SD, *n* = 3 per condition from independent experiments, each performed in triplicate. Data points represent mean number of colonies from triplicates for each independent experiment. One-way ANOVA with Tukey’s post-hoc test, (F (3, 8) = 24.6799).

In soft ECM, constitutive activation of YAP using YAP-4SA significantly increased EpH4 acinar size compared to wildtype cells (Fig. 5b). Similarly, expression of TEAD2dn was sufficient to rescue the increase in acinar size when cells were grown in stiff ECM (Fig. 5b). We also examined the effect on DNA damage by quantification of γ-H2AX foci. Expression of YAP-4SA was sufficient to cause an increase in γ-H2AX foci in cells cultured within a soft ECM, suggesting the activation of YAP signalling could promote genomic damage (Fig. 5c). However, EpH4 cells expressing TEAD2dn in stiff matrices failed to return DNA damage levels to those of wildtype cells grown in soft ECM (Fig. 5c). Furthermore, neither activation nor inhibition of YAP signalling in EpH4 cells altered the number of colonies formed when cells were seeded in soft agar following culture in 3D ECM (Fig. 5d).

These data suggest that the YAP pathway has a role in driving some of the morphological changes associated with the mechanosignalling response of MECs in 3D cultures. However, YAP signalling is not responsible for the increased DNA damage observed in cells cultured in stiff matrices and does not drive their subsequent transformation.

### Activation of RhoA signalling drives reactive aldehyde accumulation and DNA damage in cells cultured in stiff ECM

We next chose to examine the role of RhoA signalling in the DNA damage phenotype observed in cells cultured in stiff ECM. Rho signalling was a key pathway identified in the GO analysis of cells in soft and stiff 3D IPNs (Fig. 2d). Furthermore, RhoA is a key regulator of mechanical response in cells, and previous studies have implicated RhoA in driving transformation of MECs cultured in stiff 3D matrices^16^. We examined the role of RhoA signalling by generating EpH4 cell lines expressing two variants: RhoA-Q63L, which exhibits impaired GTP hydrolysis, resulting in sustained RhoA activity in response to a stimulus; and RhoA-T19N, which attenuates RhoA activity by acting as a dominant negative. Stable EpH4 lines expressing each were generated using lentiviral transduction. We then compared the effect of activating RhoA in a soft ECM and inhibiting it in a stiff ECM. We quantified acinar size, γ-H2AX foci number and colony formation in soft agar.

Similar to that observed for the YAP signalling variants, increased RhoA activity in the RhoA-Q63L cells resulted in a small but significant increase in acinar size compared to WT cells (Fig. 6a). Conversely, RhoA-T19N expressing cells formed smaller acini in stiff 3D-cultures compared to WT cells (Fig. 6a). Assessing DNA damage and transformation showed that RhoA-Q63L expression in soft conditions did not drive any significant increase in DNA damage or colony formation in soft ECM (Fig. 6b-c). However, inhibiting RhoA signalling in stiff ECM through expression of RhoA-T19N attenuated the accumulation of DNA damage, as shown via reduced number of γ-H2AX foci, and decreased colony formation in soft agar (Fig. 6b-c). Indeed, both DNA damage and colony formation were reduced to the levels observed in WT cells grown in soft 3D ECM.

**Figure 6.**
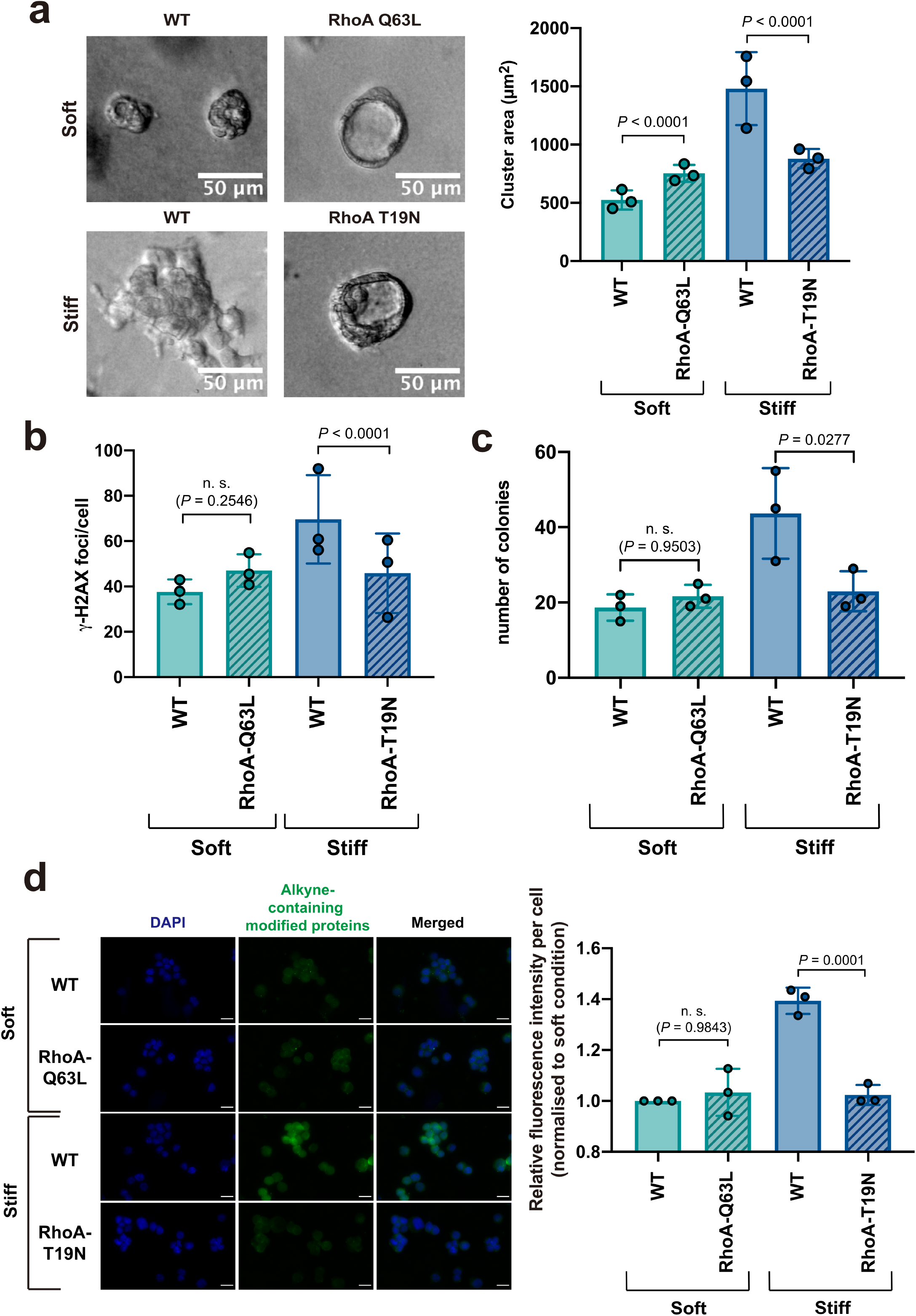
Activation of RhoA signalling drives reactive aldehyde accumulation and DNA damage in cells cultured in stiff ECM. **(a)** EpH4 cells, WT or expressing either RhoAQ63L or RhoAT19N, were grown in soft or stiff 3D ECM, as indicated. Right panel shows representative brightfield images of EpH4 acini following 10 days of culture in gels of different stiffnesses (scale bars, 50μm). Left panel shows accompanying quantification. Mean ± SD, *n* = 3 per condition from independent experiments. Data points represent mean cluster area for each independent experiment, calculated from 40-100 cells/condition. Two-way ANOVA with Tukey’s post-hoc test, (F (3, 917) = 158.3759). **(b)** Quantification of phospho-γH2AX foci (Ser139) in EpH4 cells, WT or expressing either RhoAQ63L or RhoAT19N, following 24 hours of 3D culture. Mean ± SD, *n* = 3 per condition from independent experiments. Data points represent mean number of foci for each independent experiment, calculated from 15-50 cells/condition. Two-way ANOVA with Tukey’s post-hoc test, (F (3, 366) = 16.9076). **(c)** Number of colonies formed in soft agar by EpH4 cells, WT or expressing either RhoAQ63L or RhoAT19N, following 7 days culture in either soft or stiff 3D ECM as indicated. Mean ± SD, *n* = 3 per condition from independent experiments, each performed in triplicate. Data points represent mean number of colonies from triplicates for each independent experiment. One-way ANOVA with Tukey’s post-hoc test, (F (3, 8) = 8.0291). **(d)** Representative images of EpH4 cells, WT or expressing either RhoAQ63L or RhoAT19N, following treatment and staining with Click-iT™ Lipid Peroxidation Imaging Kit after culture in gels of different stiffnesses as indicated (scale bars, 20μm); and IF quantification. Mean ± SD, *n* = 3 per condition from independent experiments. Data points represent the median fluorescence intensity for each independent experiment normalised to the median value for the soft, untreated condition (>500 cells/condition). One-way ANOVA with Tukey’s post-hoc test, (F (5, 12) = 22.4990).

We next sought to determine whether RhoA-dependent DNA damage accumulation was linked to the downregulation of ALDH activity, observed initially in the RNAseq data. As several ALDH genes were potentially downregulated, we assessed the levels of alkyne-modified protein in LAA-treated cells, which is indicative of reactive aldehyde accumulation. WT, RhoA-Q63L and RhoA-T19N EpH4 cells were cultured in soft and stiff ECM as above. Thus, we compared the single cell levels of alkyne-modified protein between WT cells and those expressing RhoA-Q63L in soft 3D ECM, and RhoA-T19N in stiff ECM (Fig. 6d). As with DNA damage and colony formation, cells expressing RhoA-Q63L did not exhibit an increase in alkyne-modified protein compared to wildtype. However, inhibition of RhoA activity through expression of RhoA-T19N reduced alkyne-modified protein levels to those observed in WT cells in soft ECM.

These results demonstrate a RhoA-dependent mechanism of DNA damage accumulation via downregulation of ALDH, and decreased clearance of reactive aldehyde species. Taken together, these findings provide a mechanism by which the increased ECM stiffness observed in women with high MD leads to the genomic damage required for breast cancer initiation.

## Discussion

Understanding the mechanistic basis for how defined risk factors promote carcinogenesis is important for mitigating their effects. High mammographic density (HMD) is the largest independent risk factor for developing breast cancer, after accounting for factors such as age and BRCA status. Indeed, some estimate that HMD may account for up to one third of breast cancers^12^. HMD is associated with increased peri-ductal stromal stiffness and thus altered mechanotransduction^13,14^ Given there is now a general understanding that many aspects of cell behaviour are governed by mechanotransduction^27^, we here examined the effects of increased altered ECM stiffness on MECs, using a mechanically-tuneable 3D hydrogel system^13^. By examining global changes in gene expression induced by increased ECM stiffness, we identified significant changes within multiple metabolic pathways. In particular, we found that decreased cellular ALDH isozyme expression contributes to accumulation of reactive aldehydes, leading to increased DNA damage and acquisition of anchorage-independent growth. Our results suggest that mechanotransduction induced changes in aldehyde metabolism leading to the acquisition of the transforming mutations are a contributing factor for breast cancer risk associated with HMD.

Mechanical cues from the ECM have previously been linked to changes in cellular metabolism^18,19^. These changes in metabolic processes in response to mechanosignalling occur through both transcriptional and post-translational regulation. Here we found global changes in metabolic pathways at the transcriptional level, coordinated through differences in ECM stiffness in 3D. Out of all the gene expression changes we identified, some of the most significant changes were in oxidoreductase pathways, with members of the ALDH isozyme family in particular being highly downregulated. The ALDH family are essential in the detoxification of endogenous reactive aldehyde species by catalysing their oxidation to carboxylic acids. Failure to remove these reactive aldehydes leaves them free to react with proteins and DNA. The reaction of these aldehydes with DNA results in the formation of base adducts which ultimately lead to DNA strand breaks during DNA replication, a major source of mutations^20^. The majority of work relating to ALDH expression centres on ALDH1A1 as a stem cell marker^28 21^. In breast cancer, ALDH1A1 has been implicated in the acquisition of drug resistance, associated with poor prognosis^29 21^. We did not see changes in ALDH1A1 expression, or other changes in genes associated with stem, progenitor, and differentiated cells within the mammary epithelial cell lineage. Instead, we provide a novel, mechanosensitive mechanism by which lowered ALDH activity results in DNA damage through increased endogenous aldehydes. Interestingly, increased DNA damage is also seen in primary tissue samples taken from women with HMD^30^. Together, this suggests that ALDH and aldehyde levels may correlate with DNA damage *in vivo*. This could provide a potential mechanism for the genomic instability which leads to the increased breast cancer risk associated with HMD. In support of this hypothesis, clinical studies found that women with HMD but without cancer showed increased levels of urinary malodialdehyde, indicating a link between altered lipid metabolism and cancer risk^31,32^.

Many signalling pathways have been described to regulate mechanosensitive cell behaviour. We found a RhoA-mediated, YAP-independent mechanism of mechanotransduction leading to an increase in reactive aldehydes and DNA damage. Although YAP activity appears to influence the altered morphology and increased size of MEC acini in stiff 3D ECM, inhibiting YAP did not alleviate the DNA damage. Studies in 2D have identified that YAP nuclear translocation is dependent on perinuclear stress fibre formation resulting in the widening of nuclear pores, and disruption of this process leads to reduced YAP nuclear localisation^33 34 35^. A recent study assessing YAP dynamics in MECs in 3D found that stress fibres failed to form in mammary acini in both soft and stiff 3D hydrogels, resulting in cytosolic localisation of YAP regardless of increasing stiffness^36^. Furthermore, both primary and immortalised breast cancer cells in 3D show a lack of YAP nuclear localisation^37^. However, some reports have shown that YAP-mediated mechanotransduction in 3D can be enhanced by oncogenic KRAS and HER2, thus increasing the sensitivity of the cells to changes in their microenvironment^38^. Taken together, this may suggest that YAP has more impact in cooperation with specific oncogenic mutations in transformed MECs, and contributes less to initiation events.

The role of Rho in mechanosignalling well-documented^39^. In the context of breast cancer, disruption of Rho signalling can reverse the transformed phenotype observed in MECs in 3D collagen hydrogels upon ECM stiffening^40^. Here we demonstrate a further role for Rho-mediated mechanosignalling in cancer initiation, where it drives the accumulation of reactive aldehyde species, resulting in DNA damage. The mechanisms by which RhoA influences aldehyde metabolism remain unknown, however, it is possible that they may occur through the influence of RhoA on transcriptional regulators. RhoA-dependent SRF transcriptional activity has recently been shown to facilitate the upregulation of glutamine metabolism in MECs expressing oncogenic Myc, suggesting an important role for RhoA signalling in breast cancer metabolism^41^. Furthermore, recent papers have established the negative regulation of SREBP1 by Rho-stimulated acto-myosin contractility leading to differential regulation of lipid synthesis pathways^42 43^. In agreement, we found that genes involved in fatty acid metabolism exhibit some of the largest fold changes in stiff compared to soft 3D ECM, providing a potential candidate for how RhoA may influence metabolism in this system. Dissecting downstream RhoA interacting partners in different 3D-hydrogel conditions may establish which signalling pathways interact differently between soft and stiff conditions^44^.

In conclusion, we demonstrate a novel, RhoA-mediated regulation of oxidoreductase pathways leading to the accumulation of reactive aldehydes in stiff 3D matrices. These results provide a potential mechanism by which the increased ECM stiffness of the periductal stroma in women with HMD drives the genetic changes required for breast cancer initiation. Therefore, these data provide a possible explanation as to how HMD confers an increase in breast cancer risk.

## Methods

### Cell culture

EpH4 cells were cultured in DMEM-F12 containing 10% FBS (v/v), containing 5 μg/mL insulin and 1% penicillin/streptomycin (v/v). For differentiation to generate β-casein, growth media was removed, and cells were washed briefly in PBS. Cells were then cultured in differentiation medium (DMEM-F12, 10% FBS (v/v), 5 μg/mL insulin, 0.5 μg/mL hydrocortisone, 1% penicillin/streptomycin (v/v) and 3 μg/mL ovine prolactin). 293T cells for lentiviral generation were cultured in DMEM supplemented with 10% FBS (v/v). Primary mouse mammary epithelial cells were obtained from 8-12-week-old ICR/FVB virgin and pregnant mice as described^45^. Briefly, tissue was harvested from pregnant mice between pregnancy day 12 and 15. Following isolation, primary cells were cultured for 48 hours in a 1:1 mix of HAMS-F12 and serum fetuin mix (20 % FBS (v/v), 1 mg/ml fetuin, F12 medium, 10 μg/ml insulin, 2 μg/ml hydrocortisone, 20 ng/ml EGF, 100 μg/ml gentamicin, 10 % PS (v/v) and 0.5 μg/ml fungizone). Following this, cells were cultured in EpH4 complete media for 24 hours, followed by EpH4 differentiation media for β-casein assays, as described above.

### Generation of rBM/Alginate hydrogels

rBM/Alginate gels were generated as described previously^15^. Briefly, 9.8 mg/mL Matrigel (Corning CAT) was combined with 25 mg/mL alginate (Pronova CAT) in a 2:1 ratio on ice. 0.5 – 1×10^5^ cells were then mixed in with each gel mixture. 50 μL 0 mM (blank DMEM-F12), 2.4 mM or 24 mM CaS0_4_ slurry was added to the gels to generate soft, medium and stiff gels respectively. This was done by placing the CaS0_4_ slurry and 200 μL Matrigel/Alginate gel into separate 1 mL syringes, connected via a female-female Luer Lock coupler (Sigma), and mixing through 4 syringe pumps, before ejecting the final gel into a 24 well plate, pre-coated with 40-50 μL of 9.8 mg/mL Matrigel. Gels were set for 30 minutes at 37°C in a humidified incubator, prior to the addition of EpH4 complete growth media. For differentiation studies, cells were cultured inside the gels for 48 hours prior to switching to EpH4 differentiation media.

### Extraction of cells from rBM/Alginate hydrogels

Assay medium was removed, and wells were rinsed with sterile, 1X PBS. 1X Trypsin-EDTA was added and pipetted to manually break the gels. Trypsin-cell mixtures were incubated for 5 minutes at 37°C. Digested mixes from replicate wells were pooled and spun at 1000 xg for 5 minutes. Cell pellets could be resuspended for RNA extractions, soft agar colony formation assays, or for cytospinning.

### Plasmids

The pEGFP-YAP construct was a gift from Dr Joe Swift. A pEGFP-YAP-4SA was then generated by introducing S61A, S109A, S164A and S381A mutations into YAP by site-directed mutagenesis. pCDH-TEAD2dn-GFP-H2B-RFP was generated by PCR amplification of the C-terminal YAP-binding domain of TEAD2 from the pCMX-Gal4-TEAD2 construct – a gift from Kunliang Guan (Addgene: 33107)^46^. This amplified TEAD fragment was then subsequently cloned into a pCDH-GFP-H2B-RFP construct. A pCDH-GFP-RhoA T19N construct was a gift from Dr Patrick Caswell. A pCDH-BFP-RhoA.T19N construct was generated from this by PCR amplification and subsequent insertion of the RhoA.T19N into a pCDH-EF1a-tagBFP construct. The pCDH-BFP-RhoA.Q63L construct was generated from the pCDH-BFP-RhoA.T19N construct through N19T and Q63L mutation of the RhoA by site-directed mutagenesis. All constructs were verified by Sanger sequencing at GATC (Eurofins).

### Lentiviral generation and transduction of EpH4 cells

HEK-293T cells with 3 μg packaging vector pMD2.G, and 4.5 μg packaging vector psPax2 using 1X PEI transfection reagent overnight. 6 μg of the plasmid containing the gene of interest was transfected alongside the packaging vectors. Transfection media was then discarded and replaced with media containing 1% (v/v) sodium butyrate (Merck-Millipore) for 6-8 hours. Cells were then returned to standard growth media for 36 hours. Following this, media containing lentivirus was harvested, and filtered through a 0.45 μM filter. Lentivirus was then precipitated through addition of 4X PEG lentivirus precipitation solution (0.05 M PEG-800, 1.2 M NaCl in 1X PBS, (pH 7.4)) and incubation for 12-72 hours at 4°C. Lentivirus was then pelleted by centrifugation at 1500 xg for 30 mins. The supernatant was then removed and centrifuged for a further 5 mins. The two pellets were then combined and resuspended in a small amount of growth media and stored at −80°C prior to transduction.

For transduction, 1×10^4^ EpH4 cells were cultured for 24 hours in EpH4 growth media. The next day, media was changed to EpH4 growth media containing 0.1% (v/v) polybrene infection/transfection reagent (Merck-Millipore), and lentivirus was added to the cells for 24 hours. Cells were then washed three times in complete growth media and once in PBS, and cultured for 2-3 weeks prior to sorting of transduced cells via FACS.

### Quantitative reverse transcription PCR (RT-qPCT

RT-qPCR gene expression analysis was performed using either TaqMan Fast Advanced Master Mix or Fast SYBR Green Master Mix according to the manufacturer’s protocols on a StepOnePlus qPCR machine (Applied Biosystems). Fluorescence was used to calculate 2^-(ΔΔCT)^ for statistical analysis using the ΔΔCT method. Values obtained for GAPDH were used to normalise values for the genes of interest. Primers used for the SYBR Green method are as follows: GAPDH: forward – 5’-GGTGAAGGTCGGAGTCAACGG-3’, 5’-GAGGTCAATGAAGGGTCATTG-3’; Sox9 (tbc); Egr2 (tbc); Snail2 (tbc). Probes used for the TaqMan method are as follows: GAPDH Mm99999915_g1; Csn Mm04207885_m1; PrlR Mm00599957_m1; Krt5 Mm01305291_g1; Krt14 Mm00516876_m1; Vim Mm01333430_m1; Elf5 Mm00468732_m1; Krt18 Mm01601704_g1; Cdh1 Mm01247357_m1. Primers for Sox9, EGR2 and Snail2 were from^47^.

### RNAseq

Polyadenylated mRNA was purified from 1 μg total RNA using poly-T oligo attached magnetic beads. Quality and integrity of total RNA was confirmed using a 2200 TapeStation (Agilent Technologies). mRNA libraries were then generated using TruSeq Stranded mRNA Assay (Illumina inc.) according to the manufacturer’s protocol. Multiplex libraries were then generated using adaptor indices and pooled prior to cluster generation using a cBot instrument. The loaded flow-cell was then paired-end sequenced (76 + 76 cycles, plus indices) on an Illumina HiSeq4000 instrument. Unmapped paired-end sequences from an Illumina HiSeq2500 sequencer were tested by FastQC, (http://www.bioinformatics.babraham.ac.uk/projects/fastqc/). Sequence adapters were removed, and reads were quality trimmed using Trimmomatic^48^. The reads were mapped against the reference mouse genome (mm10/GRCm38) and counts per gene were calculated using annotation from GENCODE M2 (http://www.gencodegenes.org/) using Tophat2^49^ and HTSeq^50^. Normalisation, Principal Componenents Analysis, and differential expression was calculated with DESeq2^51^.

### Gene Ontology Analysis

Over-represented GO terms of significantly changing genes (adjusted p< 0.05 and fold change ±2) were calculated by the R package TopGO^52^. GO terms with adjusted p-values < 0.05 were considered significant^53^ For the GO-gene bipartite graph, the significantly changing genes were submitted for to GO overrepresentation analysis using the R package ClusterProfiler^54^ ClusterProfiler results were simplified using semantic similarity^55^ to cluster terms with similarity score > 0.7, with the representative term taken from each cluster with the lowest adjusted p-value. GO terms with adjusted p-values < 0.05 were considered significant.

### Immunofluorescence staining

Cells isolated from gels were resuspended in 1X PBS, and 50 μL was cytospun at 400 RPM for 5 minutes onto polysine slides using a Shandon Cytospin 2 (Thermo Scientific). Cells were then fixed onto the slides with 4% PFA. Fixed cells were permeabilized using 0.5 % Triton-X for 20 minutes, then washed in 1X TBS. The cells were then blocked using 3 % BSA in TBS for at least 1 hour before addition of the primary antibody in antibody dilution buffer (1% horse serum (v/v), 0.05% Tween 20 (v/v), 0.2% Triton-X-100 (v/v), and 0.05% sodium azide (v/v) in 1X PBS. Cells were incubated with primary antibody for at least 1 hour at room temperature, or overnight at 4°C. The primary antibody solution was then removed and cells washed several times with 1X TBS before addition of the secondary antibody in antibody dilution buffer. Cells were incubated with secondary antibody solution for 1 hour at room temperature, then washed several times with 1X TBS. Cells were then incubated with DAPI (0.1 μg/ml) for 10 minutes, then washed several times in 1X TBS and once in ddH_2_0. Cytospun cells were covered with a coverslip and DAKO fluorescent mounting medium (Agilent Technologies). Staining was visualised using a Imager.M2 fluorescence microscope (Zeiss). Images were taken using a Hamamatsu ORCA-ER Digital Camera.

### Lipid peroxidation

WT EpH4 cells, and EpH4 cells expressing BFP-RhoA-Q63L or BFP-RhoA-Q63L were grown in soft and stiff rBM/Alginate gels for 24hrs. 5 hours prior to extraction from the gels, 50 μM LAA was added to the media. Cells were then fixed in 4% PFA, washed once in PBS, and cytospun onto polysine slides. Staining from lipid peroxidation was then carried out using a Click-iT Lipid Peroxidation Imaging Kit – Alexa Fluor 488 (Invitrogen) according to the manufacturer’s protocol and mounted with DAKO fluorescent mounting medium (Agilent Technologies). Images were acquired on a 3D-Histech Pannoramic-250 microscope slidescanner using a 40x/0.95 Plan Apochromat objective (Zeiss) and the DAPI and FITC filter sets. Quantification of fluorescence intensity was completed using QuPath (0.2.0-m8)^56^ and images were exported and processed in Fiji^57^.

### Colony-formation assay

1 mL of molten 0.8% low gelling agarose solution was added to the bottom of each well of a 6 well plate and allowed to set for 20 minutes at room temperature. Following this, EpH4 cells grown for 7 days in rBM/Alginate gels were extracted and 1×10^4^ seeded inside 0.4% low gelling agarose solution and cast on top of the base layer. Plates were then placed on ice to cool until set. 1 mL of growth media was then added to each well and cells were incubated for 21 days at 37°C and 5% CO_2_ under humidified conditions, with media replaced every 4 days. Following incubation, cells were fixed in 0.005% crystal violet solution in 4% PFA for 1 hour. Number of colonies were then counted manually counted using a Leica DMIL LED Inverted Brightfield microscope.

### Atomic force microscopy

Samples were measured in water using a Bioscope Catalyst AFM mounted on a Nikon Eclipse Ti inverted microscope with a silicon nitride MLCT probe (Microlever series; E; radius 20 nm). Spring constant was maintained at 0.149 Nm-1. Nanoscope Software v 8.15, Bruker was used to collect data.

### Statistical Analysis

All statistical analyses were carried out using PRISM 8 for MacOS (version 8.4.3 (471)). Where appropriate, statistical significance was determined by Two-tailed student’s *t* test, one- or 2-way ANOVA with Tukey’s post-hoc test, or Kruskal Wallis test with Dunn’s multiple comparisons post-hoc test. The test used for each experiment is defined in the figure legends.

## Acknowledgements

AW was supported by an MRC doctoral training award. HS was a recipient of a University of Manchester Presidents Doctoral Scholarship. MJ and HP were funded through the Manchester Cancer Research Centre by CRUK training awards. JS was funded by a Biotechnology and Biological Sciences Research Council (BBSRC, UK) David Phillips Fellowship (BB/L024551/1). The Wellcome Centre for Cell-Matrix Research is supported by a core grant from the Wellcome Trust (203128/Z/16/Z). We are grateful for the assistance provided by staff in the Bioimaging, Bio-MS core facilities.

## Author contributions

AW, HS KB and APG conceived and planned the project. AW, HS, MJ and HP conducted the majority of the experiments presented in the paper. EB established the lipid peroxidation assay. EZ and CL performed image and bioinformatics analysis respectively. MJ, HP, JS, KB and APG wrote the paper. All authors discussed the results and commented on the manuscript.

