## Supplementary figures 1-3 for "Increased microenvironment stiffness leads to altered aldehyde metabolism and DNA damage in mammary epithelial cells through a RhoA-dependent mechanism"

Supplementary data.

**Supplementary figure legends.**

**Figure S1. (a)** Schematic of the mechanically tuneable, 3D Matrigel/alginate culture model (left panel). The addition of increasing concentrations of $CaSO_4$ results in increasingly stiffer gels, as measured by atomic force microscopy (AFM) (right panel), (n = 3).

**(b)** Representative brightfield images of EpH4 acini following 10 days of culture in gels of different stiffnesses (scale bars, 100μm), and accompanying quantification. Mean ± SD, *n* = 3 per condition from independent experiments, each performed in duplicate. Data points represent mean cluster area from duplicates for each independent experiment. Two-way ANOVA with Tukey's post-hoc test, (F (2,9) = 20.6639).

**Figure S2. (a)** Hierarchical clustering heatmap representing significantly differentially expressed genes in EpH4 cells cultured in 2D and 3D gels of different stiffnesses, as determined by RNAseq. **(b)** Volcano plot generated from RNAseq data, showing genes that are significantly upregulated (blue) and downregulated (red) in EpH4 cells grown in the stiff condition, relative to soft. Upper quadrant represents 3D cultures, and lower quadrant respresents 2D cultures.

**Figure S3.** $Log_2$ fold-change in expression of genes associated with the major DNA damage repair pathways in EpH4 cells. Expression changes are shown in the stiff condition relative to soft, as determined by RNAseq data. Pathways are: **(a)** base excision repair; **(b)** nucleotide excision repair; **(c)** mismatch repair; **(d)** homologous recombination; and **(e)** non-homologous end joining. Error bars represent SE (*n* = 3), statistical significance was determined using DESeq2.

**Figure S4. (a)** Quantification of phospho-CHK1 (Ser345) and phospho-CHK2 (Thr68) in EpH4 cells cultured in soft or stiff 3D ECM for 24 hrs. Mean ± SD, *n* = 2 per condition from independent experiments, each performed in duplicate. Data points represent mean number of foci for each independent experiment, calculated from 20-30 cells/condition. Two-way ANOVA, (F (1,100) = 29.4201) and (F (1,91) = 27.5400) for phospho-CHK1 and phospho-CHK2, respectively.

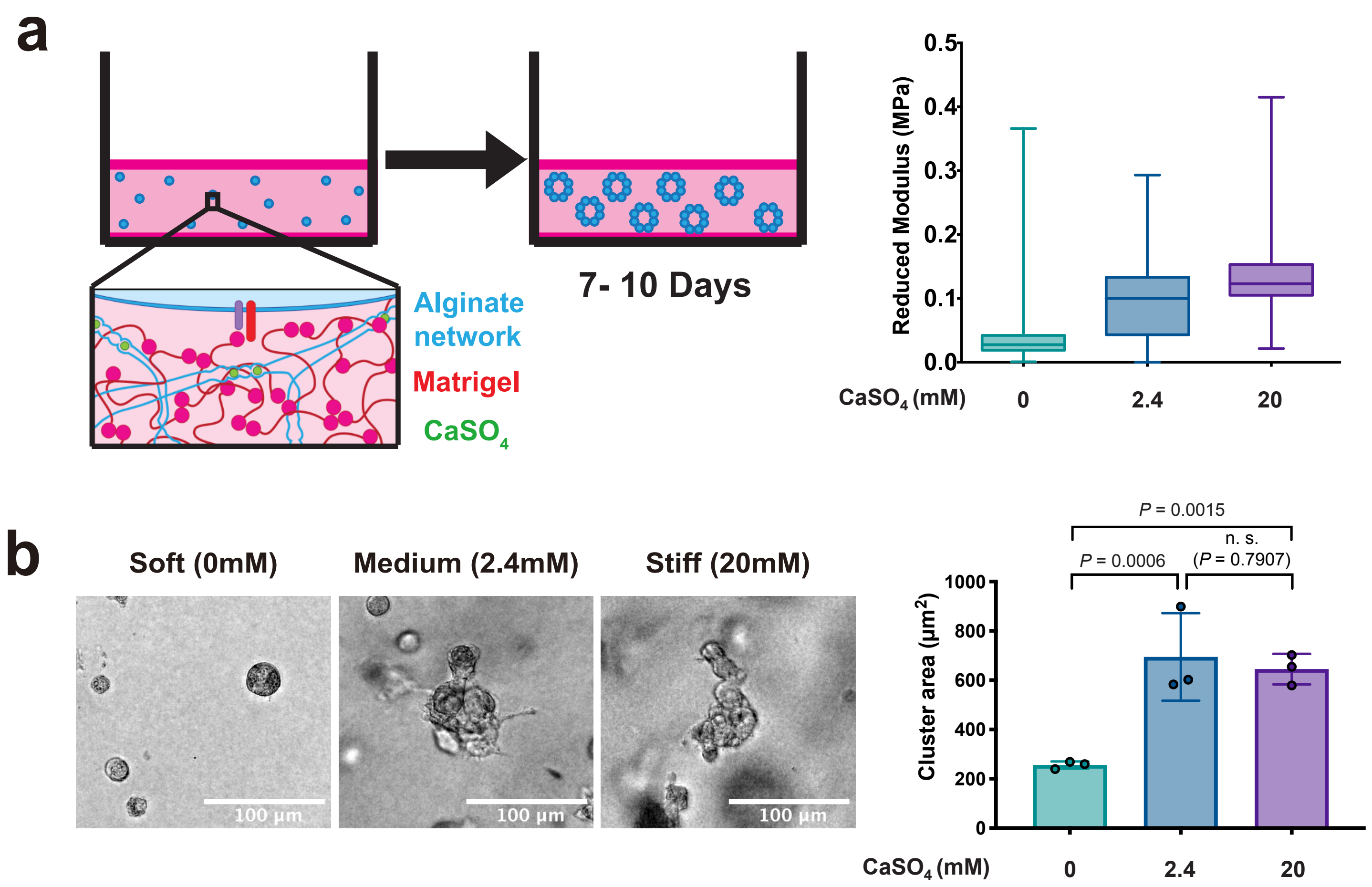
a
7- 10 Days
Alginate network
Matrigel
CaSO4
Reduced Modulus (MPa)
0.5
0.4
0.3
0.2
0.1
0.0
CaSO4 (mM)
0
2.4
20
b
Soft (0mM)
Medium (2.4mM)
Stiff (20mM)
100 µm
100 µm
100 µm
P = 0.0015
n. s.
P = 0.0006
(P = 0.7907)
Cluster area (µm2)
1000
800
600
400
200
0
CaSO4 (mM)
0
2.4
20

Figure S1

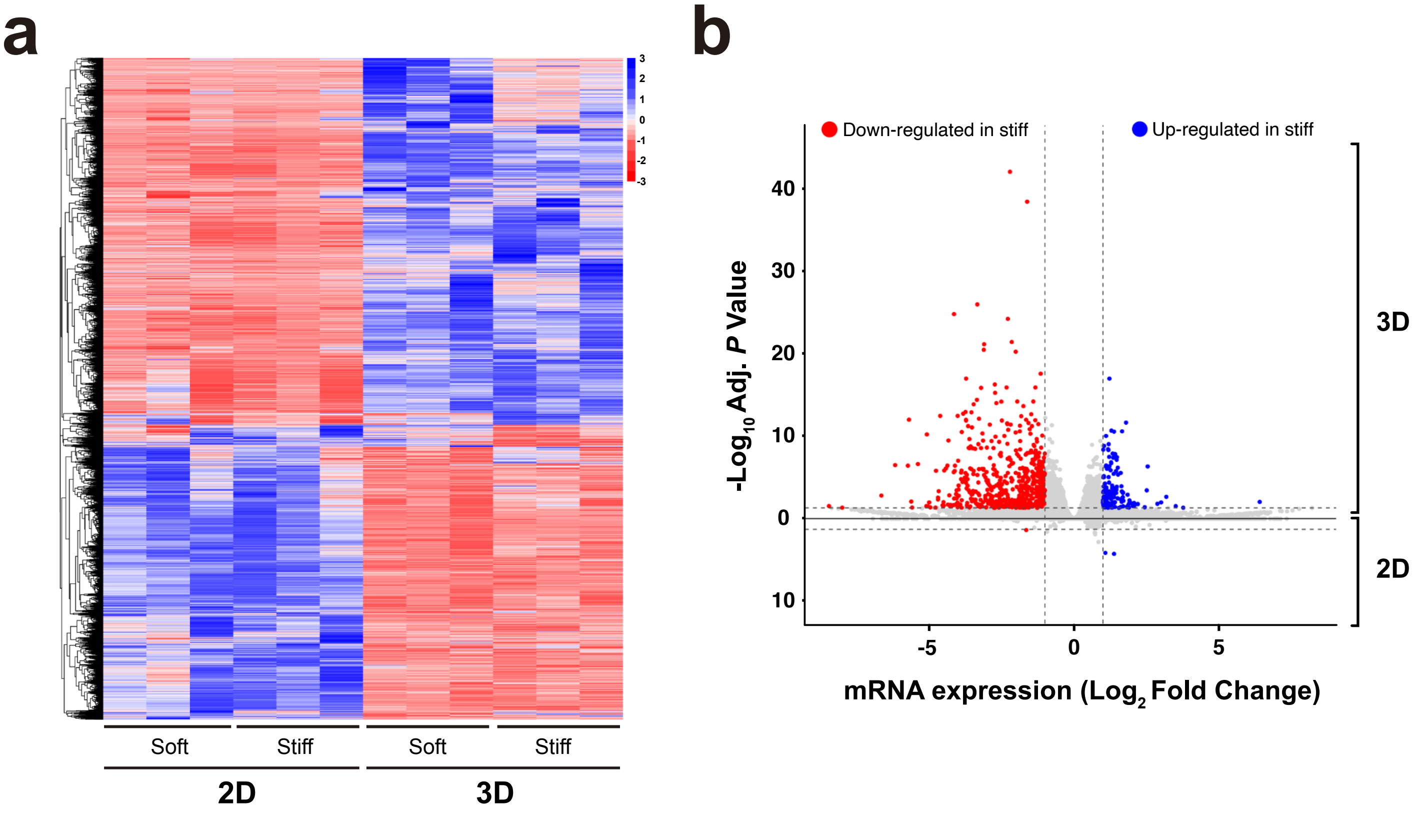
a
3
2
1
0
-1
-2
-3
Soft
Stiff
Soft
Stiff
2D
3D
b
Down-regulated in stiff
Up-regulated in stiff
-Log10 Adj. P Value
40
30
20
10
0
10
3D
2D
-5
0
5
mRNA expression (Log2 Fold Change)

Figure S2

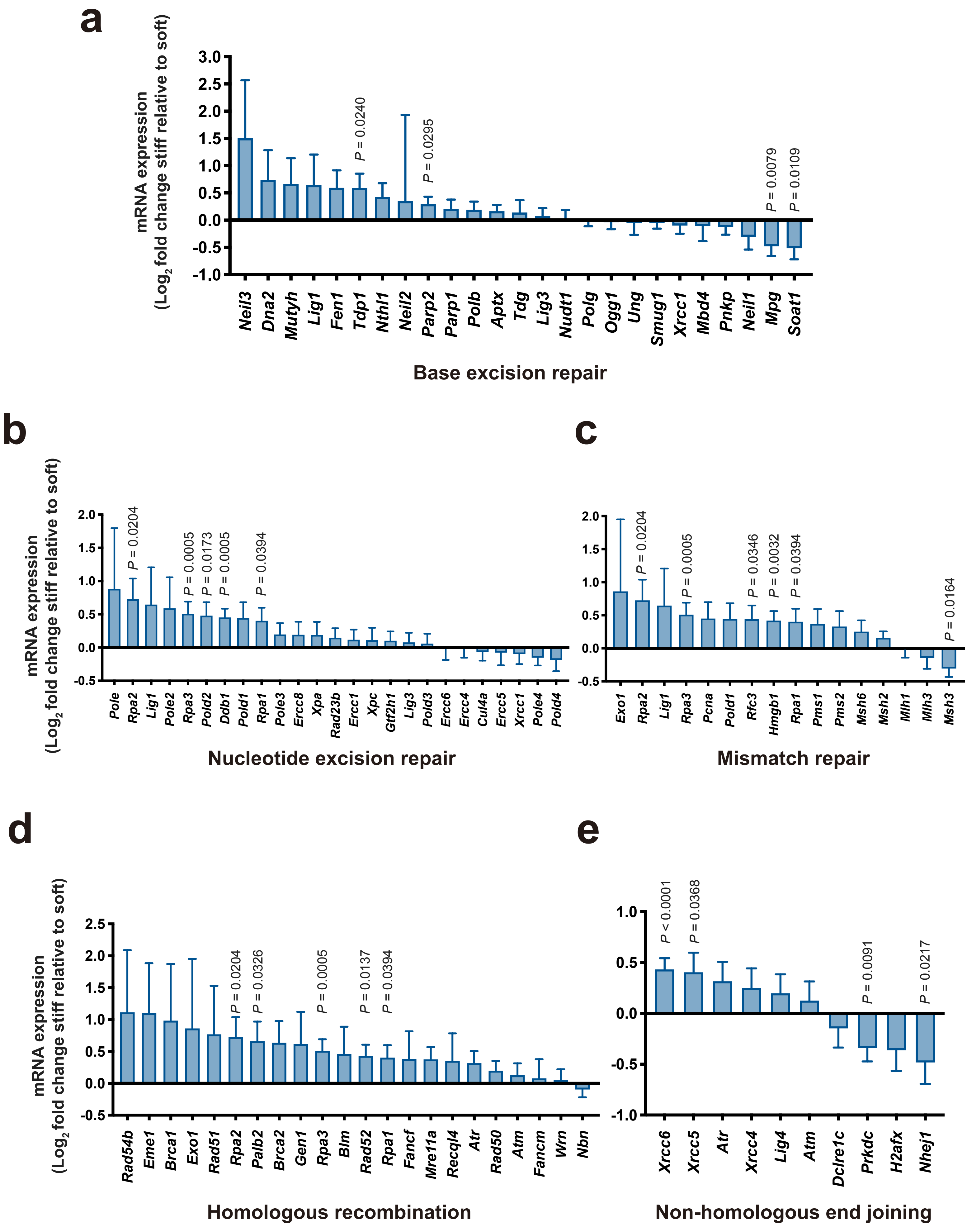

a
mRNA expression
(Log2 fold change stiff relative to soft)
3.0
2.5
2.0
1.5
1.0
0.5
0.0
-0.5
-1.0
P = 0.0240
P = 0.0295
P = 0.0079
P = 0.0109
Neil3
Dna2
Mutyh
Lig1
Fen1
Tdp1
Nthl1
Neil2
Parp2
Parp1
Polb
Aptx
Tdg
Lig3
Nudt1
Polg
Ogg1
Ung
Smug1
Xrcc1
Mbd4
Pnkp
Neil1
Mpg
Soat1
Base excision repair
b
mRNA expression
(Log2 fold change stiff relative to soft)
2.0
1.5
1.0
0.5
0.0
-0.5
P = 0.0204
P = 0.0005
P = 0.0173
P = 0.0005
P = 0.0394
Pole
Rpa2
Lig1
Pole2
Rpa3
Pold2
Ddb1
Pold1
Rpa1
Pole3
Ercc8
Xpa
Rad23b
Ercc1
Xpc
Gtf2h1
Lig3
Pold3
Ercc6
Ercc4
Cul4a
Ercc5
Xrcc1
Pole4
Pold4
Nucleotide excision repair
c
2.0
1.5
1.0
0.5
0.0
-0.5
P = 0.0204
P = 0.0005
P = 0.0346
P = 0.0032
P = 0.0394
P = 0.0164
Exo1
Rpa2
Lig1
Rpa3
Pcna
Pold1
Rfc3
Hmgb1
Rpa1
Pms1
Pms2
Msh6
Msh2
Mlh1
Mlh3
Msh3
Mismatch repair
d
mRNA expression
(Log2 fold change stiff relative to soft)
2.5
2.0
1.5
1.0
0.5
0.0
-0.5
P = 0.0204
P = 0.0326
P = 0.0005
P = 0.0137
P = 0.0394
Rad54b
Eme1
Brca1
Exo1
Rad51
Rpa2
Palb2
Brca2
Gen1
Rpa3
Blm
Rad52
Rpa1
Fancf
Mre11a
Recql4
Atr
Rad50
Atm
Fancm
Wrn
Nbn
Homologous recombination
e
1.0
0.5
0.0
-0.5
-1.0
P < 0.0001
P = 0.0368
P = 0.0091
P = 0.0217
Xrcc6
Xrcc5
Atr
Xrcc4
Lig4
Atm
Dclre1c
Prkdc
H2afx
Nhej1
Non-homologous end joining


**Figure S3**

a

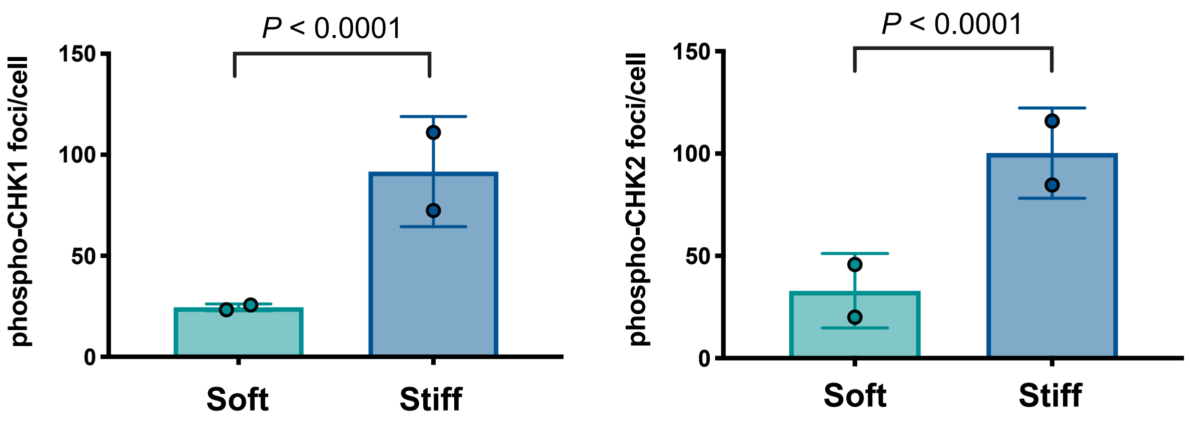
P < 0.0001
150
100
50
0
phospho-CHK1 foci/cell
Soft
Stiff
P < 0.0001
150
100
50
0
phospho-CHK2 foci/cell
Soft
Stiff

Figure S4
